# Targeted RNA-seq improves efficiency, resolution, and accuracy of allele specific expression for human term placentas

**DOI:** 10.1101/2021.01.25.428155

**Authors:** Weisheng Wu, Jennie L. Lovett, Kerby Shedden, Beverly I. Strassmann, Claudius Vincenz

## Abstract

Genomic imprinting is an epigenetic mechanism that results in allele specific expression (ASE) based on parent of origin. It is known to play a role in the prenatal and postnatal allocation of maternal resources in mammals. ASE detected by whole transcriptome RNA-seq (wht-RNAseq) has been widely used to analyze imprinted genes using reciprocal crosses in mice to generate large numbers of informative SNPs. Studies in humans are more challenging due to the paucity of SNPs and the poor preservation of RNA in term placentas and other tissues. Targeted RNA-seq (tar-RNAseq) can potentially mitigate these challenges by focusing sequencing resources on the regions of interest in the transcriptome. Here we compared tar-RNAseq and wht-RNAseq in a study of ASE in known imprinted genes in placental tissue collected from a healthy human cohort in Mali, West Africa. As expected, tar-RNAseq substantially improved the coverage of SNPs. Compared to wht-RNAseq, tar-RNAseq produced on average four times more SNPs in twice as many genes per sample and read depth at the SNPs increased 4-fold. In previous research on humans, discordant ASE values for SNPs of the same gene have limited the ability to accurately quantify ASE. We show that tar-RNAseq reduces this limitation as it unexpectedly increased the concordance of ASE between SNPs of the same gene, even in cases of degraded RNA. Studies aimed at discovering associations between individual variation in ASE and phenotypes in mammals and flowering plants will benefit from the improved power and accuracy of tar-RNAseq.

## Introduction

Genomic imprinting is an epigenetic phenomenon that results in allele specific expression (ASE) based on parent of origin. Many imprinted genes are found in the nutritive tissue of placental and marsupial mammals as well as flowering plants^1,2^. Under the kinship hypothesis, genomic imprinting evolved due to a conflict of interest between the genes an offspring inherited from its mother versus its father over the amount of resources to be allocated to the current offspring^3^. One consequence of imprinting is that, for specific genomic regions, the paternal and maternal genomes are not equivalent. Evidence that both are required for normal development derives from a series of mouse studies in the 1980s that generated gynogenotes or androgenotes. In uniparental disomies (UPD), non-equivalency was limited to certain genomic regions that later were identified as clusters of imprinted genes^1^. In humans, about 100 imprinted genes have been identified^4,5^. The highest proportion of imprinted genes were expressed in embryonic, extra-embryonic and brain tissues^5^, and impacted neurological development, placentation and fetal growth^6^. Regulation of imprinting is governed by imprinting control regions (ICRs) through epigenetic mechanisms involving DNA methylation, lncRNAs, histone modifications, and high-order chromatin organization^7,8^.

High throughput sequencing technologies including RNA-seq and DNA methylation sequencing have been widely used to study genomic imprinting^9^. Transcriptome wide ASE is determined by combining quantification of whole transcriptome RNA-seq (wht-RNAseq) reads with identification of heterozygous SNPs in DNA^10^. Animal studies gain additional power from reciprocal cross breeding of closely related strains, which produces higher SNP densities and phased reference genomes that pinpoint the parent of origin of each allele at every SNP. These crosses permit imprinting to be distinguished from sequence dependent allelic expression bias^5,11–13^. In humans, fewer SNPs are present than in crossbred animal models due to lower genetic diversity; nonetheless RNA-seq has been successfully employed in many human tissues^14–19^. Frequently, the parent’s genotype is not available in human studies and ASE is determined without the parent of origin of the bias^4,5,20^.

To quantify ASE from RNA-seq, best practice protocols have been proposed to accommodate several technical factors^21^. For example, appropriate alignment methods should be used to reduce the tendency for mapping bias to favor the reference alleles^22,23^. A few studies showed that the accuracy of ASE quantified from RNA-seq was especially limited when the read depth on the measured SNPs was insufficient^24–26^, which could lead to low power to predict imprinting and poor agreement of ASE between the SNPs from the same genes^27^ and even from the same exons^28^.

The consensus is that most imprinted genes in humans and mice have been identified with these genome wide approaches^4,5,14^. However, the well documented population variability in imprinting and potential phenotypic effects are poorly understood^14,29^. Thus, there is a need to quantify ASE with high precision in a population setting in a cost-efficient manner. In a previous study we used wht-RNAseq to measure ASE in 91 known imprinted genes in human term placentas collected from a cohort in Mali. We showed that departures from mono-allelic RNA expression were prevalent in many imprinted genes in this cohort. The number of reads we obtained from imprinted genes was limited because many highly expressed genes in placenta are not imprinted and constituted a large fraction of the total reads^29^.

To overcome this limitation, we employed a tar-RNAseq approach to focus sequencing resources on the genes of interest. We enriched RNA against a targeting panel designed to cover exonic regions of 520 genes and quantified ASE on the informative SNPs in this panel. We hypothesized that this tar-RNAseq dataset could substantially increase the coverage of SNPs and genes of interest and improve the accuracy of ASE determination, compared to our previously published wht-RNAseq dataset. In order to test this hypothesis, we performed the same ASE analysis and compared the results for genes common to both datasets. For 75 genes, reported in the literature to be imprinted, we had at least one well covered heterozygous SNP in both our tar-RNAseq and wht-RNAseq datasets. In support of our hypothesis, we show that tar-RNAseq covered many more informative SNPs and greatly improved the SNP read depth, which allowed us to measure ASE at more sites in more genes. We also obtained two results that were not expected by the deeper sequencing of a targeted approach. First, tar-RNAseq produced much higher concordance of ASE between the SNPs from the same genes resulting in improved quantification of the gene-level expression bias. Second, tar-RNAseq in combination with rRNA depletion permitted efficient ASE determination from degraded RNA whereas higher RNA integrity was required for wht-RNAseq. These improvements have enabled us to quantify the inter-individual variability of ASE in our cohort with high resolution and accuracy, which will be critical for querying associations between genomic imprinting and growth phenotypes. Our results provide performance metrics for this approach on samples collected in the field, which can be applied to design ASE studies in other populations or species.

## Methods

### IRB

Informed consent or assent was obtained from participants depending on whether they were adults or children. IRB approval was obtained from the University of Michigan IRBMED (HUM00043670) and from La Faculté de Médecine de Pharmacie et d’Odontostomatologie (FMPOS) de Bamako in Mali (N°2016/68/CD/FMPOS).

### Capture design

Sample collection, nucleic acid purification, and wht-RNAseq were described previously^29^. The capture region for tar-RNAseq included exonic regions for all genes with reports in the literature indicating imprinted expression or allelic methylation. The criteria for inclusion were non stringent to avoid the exclusion of imprinted genes at the cost of including some non-imprinted genes. Furthermore, the targeted genes included genes relevant to diseases that are of interest in this cohort (n=67), and genes with consistent high placental expression in the wht-RNAseq dataset (n=71, Supplementary Table 1). ERCC spike-in controls were also targeted except for nine transcripts that spanned the expression range. Genetic variation identified in the cohort was taken into account by including 100 bp capturing oligonucleotides containing the Alt allele for all SNPs spaced more than 50 bp apart (n=4634). The targeting regions were evaluated by the NimbleDesign Software and oligonucleotides covering 2,797,406 bases were synthesized using the Roche SeqCap RNA Developer platform (Supplementary .bed file).

### Genotyping

Genotyping of F2 umbilical cord tissues (n = 227) and F1 saliva samples (n = 189) was performed with targeted DNA sequencing (Roche: SeqCap EZ Choice). The region genotyped for the tar-RNAseq samples overlapped the region of the wht-RNAseq samples (1.4 Mb) and included more genes for a total of 3.9 Mb. The analysis presented here is limited to the 75 genes that had RNAseq data in both datasets. This subset of genes mapped to 0.58 Mb and 0.48 Mb of the regions genotyped in the tar-RNAseq and the wht-RNAseq samples, respectively. Library preparation and hybridization captures were performed at the University of Michigan Advanced Genomics Core following manufacturer’s protocols.

### Tar-RNAseq Library preparation and sequencing

The University of Michigan Advanced Genomics Core prepared KAPA RNA HyperPrep Kit (Roche KK8540) libraries or KAPA RNA HyperPrep with RiboErase Kit (Roche KK8560) libraries from 1000 ng, DNaseI digested, placental total RNA using conditions adapted to each sample’s RNA quality. Initially, only samples with poorly defined rRNA bands on Agilent traces (RIN <2.5) were depleted of rRNA with RiboErase prior to fragmentation. Later, samples with intermediate RNA (RIN <6.0) were also processed with RiboErase as the cost was not prohibitive. Fragmentation conditions were established based on each sample’s Agilent Bioanalyzer DV_200_ quality metric which reflects the percentage of RNA fragments above 200 nucleotides: DV_200_ < 55 at 65 °C for 1 min; 55 > DV_200_ <70 at 65 °C for 4 min; DV_200_ > 70 and RIN < 3.8 at 85 °C for 4 min; DV_200_ > 70 and RIN >3.8 94 °C for 4 min. ERCC exogenous RNA controls (ThermoFisher Scientific 4456739) were included in all library preparations according to manufacturer’s guidelines. Six indexed cDNA libraries were pooled for each capture reaction totaling 1 μg of cDNA. In cases where Kapa RNA HyperPrep plus RiboErase libraries were multiplexed with Kapa RNA HyperPrep (non-rRNA-depleted) libraries, the amount of the rRNA-depleted library was adjusted to10-fold less than non-depleted RNA libraries in these mixtures. Libraries were sequenced on an Illumina NovaSeq (S4). RNAs with RIN < 3.8 and DV_200_ < 70 were generally selected for KAPA RNA Prep Plus RiboErase library preparations. In total, 236 RNA samples from 227 F2 individuals were sequenced.

### Pyrosequencing

Allelic expression of select heterozygous SNPs was validated by pyrosequencing. cDNA synthesis by RT was performed immediately after DNaseI digestion of placental RNA with the ProtoScript^®^II First Strand cDNA Synthesis Kit (E6560, New England Biolabs) and random hexamer primers. Qiagen PyroMark Assay Design 2.0 software was used to design pyrosequencing primers and amplicons were generated with PyroMark PCR Kit (978705, Qiagen) and sequenced using a PyroMark Q96 MD workstation. 19 SNPs were pyrosequenced in 3 placentas. The Pearson correlation coefficient for major allele frequency between RNAseq and pyrosequencing was 0.98 (p=9.6×10^−19^).

### DNA sequencing analysis

Illumina adapter contamination and read ends with base quality < 20 were removed using Trimmomatic^30^. Reads shorter than 36 nt after trimming were discarded. Trimmed reads were aligned to hg38 reference genome using BWA^31^. Read deduping and base quality scale recalibration were performed using MarkDuplicates and BaseRecalibrator, respectively, from GATK^32,33^. SNPs and short INDELs were called using HaplotypeCaller, GenomicsDBImport and GenotypeGVCFs from GATK. Resulting variants underwent GATK-recommended hard-filtering for SNPs and INDELs separately. Furthermore, we applied a series of filters in order to remove less-confident genotypes that included the following: 1) variants with genotyping quality < 20 or total read depth < 20; 2) variants falling in the regions with 100mer-alignability score < 1 using the Umap multi-read mappability track^34^; 3) variants falling in the ENCODE Blacklist regions^35^ or the genomic SuperDups regions^36^; 4) variants with known alternate allele mapping bias identified in a previous study^21,37^; 5) variants that had more than one alternate allele; 6) heterozygous SNPs whose reference allele frequency was < 0.2 or > 0.8; 7) homozygous SNPs whose reference allele frequency was >0.05; 8) homozygous reference sites whose reference allele frequency was < 0.95; 9) SNPs where > 5% of reads supported an allele that was neither reference nor alternate; 10) SNPs exhibiting excess heterozygosity (GATK-calculated metrics ExcessHet > 54.69); and 11) SNPs having a nearby INDEL within 150bp. PhaseByTransmission in GATK was used to phase the variants in a subset of the samples (45%) where both parents were genotyped. The phased variants were filtered by requiring the transmission probability score to be no lower than 20, and then combined with the variants phased by HaplotypeCaller. Eight F2 samples were excluded from the final phasing results due to excessive Mendelian violations indicative of nonpaternity or tube error.

### RNA sequencing analysis

Previously published wht-RNAseq data^29^ was used with the allelic read counts recalculated using deduped alignments, and the same workflow was also used for the analysis of the tar-RNAseq data. Illumina adapter contamination and read ends with base quality < 20 were removed using Trimmomatic. Reads shorter than 36 nt after trimming were discarded. HISAT2^38^ was used to first build a new reference for each individual to incorporate the genomic variants identified from the corresponding DNA sample, and second to align the paired trimmed reads onto this reference with splice sites from GENCODE GTF^39^. Alignments were filtered and deduped using WASP^40^ to reduce biases. Properly paired alignments with the highest mapping quality were selected as confident alignments and used for downstream analyses.

StringTie^41^ was used to quantify the relative expression at the transcript level. Alignments were split into sense-strand and antisense-strand alignments. ASEReadCounter from GATK was used to calculate allele-specific RNA read depth in both strands at each heterozygous SNP of the paired DNA sample. SNPs were annotated with the coordinates of the exons to which they mapped and overlapping exons in the same gene were merged into one interval. SNPs covered by at least 10 reads and mapped to unique genes and transcripts expressed at > 0.1 TPM in a placental reference RNA-seq data set were retained^42^. The SNP level imprinting codes were generated after considering all genes affected by the SNP using VEP, Variant Effect Predictor^43^. Targeted enrichment was measured as one minus the off-target aligned base ratio computed by CollectHsMetrics in GATK. Maternal contamination was assessed and removed as previously described^29^.

For the comparison between tar-RNAseq and wht-RNAseq, we only used the SNPs in the genes that carried at least one SNP in at least one sample in both datasets and only paternally expressed (PEGs), maternally expressed (MEGs), and complexly expressed (CEGs) genes were considered, which limited the comparison to 75 genes (Supplementary Table 1, column D). SNP-level Pat-Freq was calculated as the ratio of the paternal allele read count to the total read count. For gene-level Pat-Freq, we summed the paternal allele read counts and total read counts from all the SNPs of the gene and calculated their ratio. To determine the ASE correlations between SNPs at the gene or exon level, Pearson correlation coefficients were calculated across all pairwise combinations of SNPs mapping to the same gene or exon.

### RNAseq library preparation cost evaluation

A comparison between the cost of wht-RNAseq and tar-RNAseq (Supplementary Table 3) on a per sample basis was made to weigh increases in sequencing depth and coverage against costs of adding a target capture step to the library preparation method. Library preparation service costs reflect pricing as of May 7, 2020 at the University of Michigan Advanced Genomics Core. All other reagents reflect pricing at the time of purchase. The custom Roche SeqCap Target Enrichment System employed for tar-RNAseq has been discontinued but similar products are currently offered by multiple suppliers (e.g. agilent.com, arborbiosci.com, illumina.com, idtdna.com, qiagen.com, thermofisher.com, twistbioscience.com)

## Results

### Congruent ASE values from Targeted RNA-seq and whole transcriptome RNA-seq

We compared allelic count distributions in tar-RNAseq and wht-RNAseq datasets for a set of 75 genes that have been reported to be imprinted in the literature and that contained at least one SNP in both datasets. The SNPs in these datasets were derived from 227 and 40 term placentas in tar-RNAseq and wht-RNAseq, respectively. For both datasets, exonic SNPs were identified by DNA-seq of umbilical cord tissue. The total size of the regions genotyped for the tar-RNAseq samples was 3.9 Mb yielding 3,647 exonic SNPs, and for the wht-RNAseq samples was 1.4 Mb yielding 2,517 exonic SNPs that mapped to the 75 genes of interest (Methods). SNPs were annotated as in our earlier study^29^ as PEGs or MEGs based on the parental bias reported in the literature, and as CEGs, for genes with complex imprinting patterns or conflicting literature data.

To verify the parent of origin for the expression bias, we phased the SNPs in the subset of samples for which parental genotypes were known and calculated paternal allele frequency (Pat-Freq) (Figure 1A). By combining transmission-based and read-based phasing, we were able to phase on average 265 and 68 SNPs per sample in 110 and 28 samples from tar-RNAseq and wht-RNAseq, respectively. In both datasets, the distributions of Pat-Freq in all three categories agreed with the previously reported imprinting directions (Figure 1A). The interquartile range of Pat-Freq in CEGs was smaller for tar-RNAseq than for wht-RNAseq. To determine allelic bias for all data, phased and unphased, we calculated ASE as |0.5 – (Reference reads/Total reads)| (Figure 1B) ^21^. Both tar-RNAseq and wht-RNAseq data showed the strongest allelic bias for PEGs, reduced allelic bias for MEGs, and close to biallelic expression for many CEGs. The ASE distribution in the tar-RNAseq data exhibited more biallelic expression in all groups (Figure 1B).

**Figure 1.**
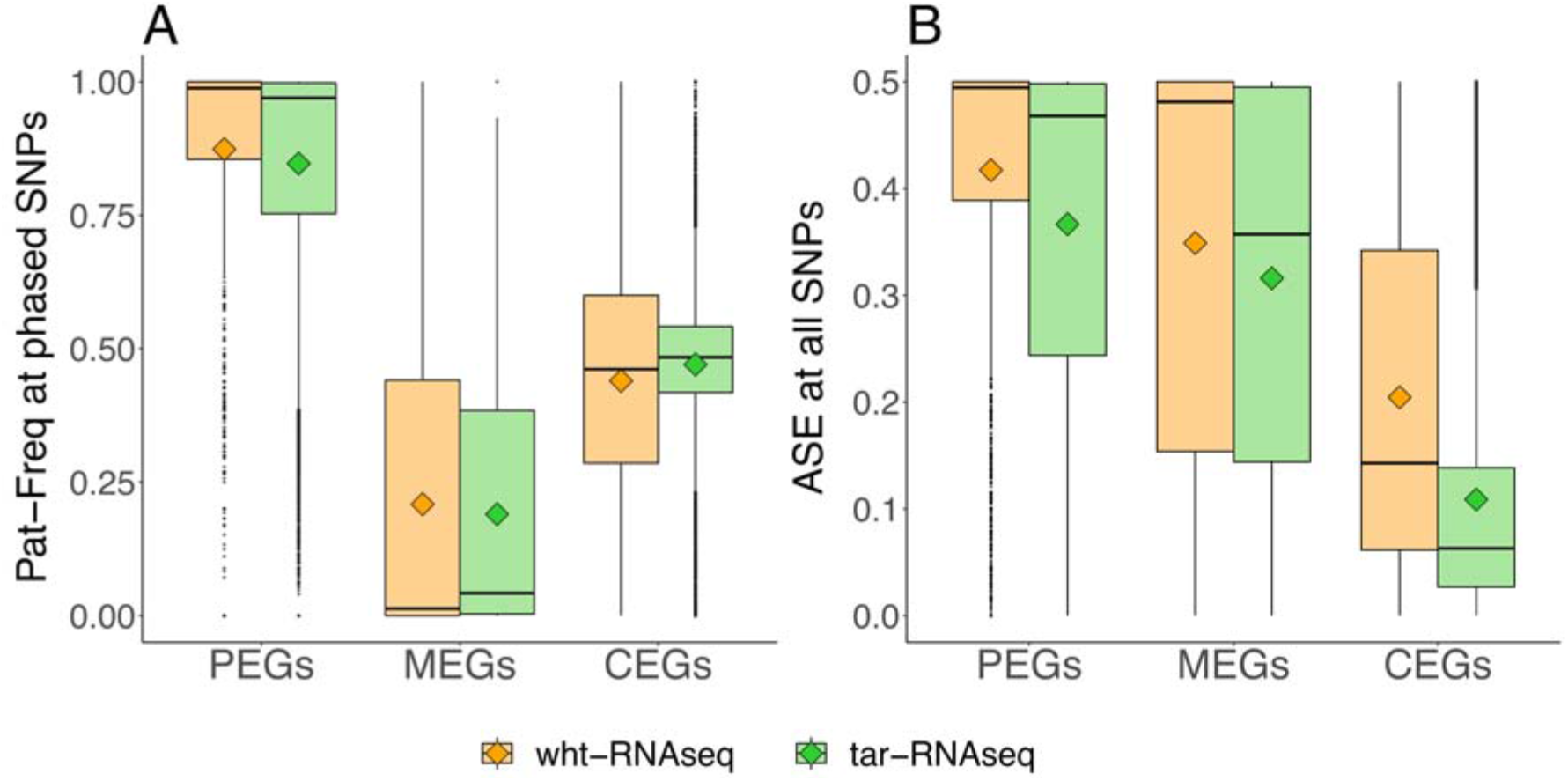
Distribution of parent of origin expression and ASE in PEGs, MEGs and CEGs. The box plots show the distributions of (a) Pat-Freq and (b) ASE at the SNPs in paternally expressed genes (PEGs), maternally expressed genes (MEGs) or complex expressed genes (CEGs). The mean of each distribution is indicated by a diamond. Orange and green colors denote wht-RNAseq and tar-RNAseq, respectively.

The agreement between datasets is further illustrated by the correlation of gene-level Pat-Freq between tar-RNAseq and wht-RNAseq data for the placenta that was assayed both ways (Figure 2). The mean Pearson correlation coefficient was 0.95 overall, 0.90 for PEGs, 1.00 for MEGs, and 0.58 for CEGs. Thus, the correlation between the two data sets for gene-level Pat-Freq was strong.

**Figure 2.**
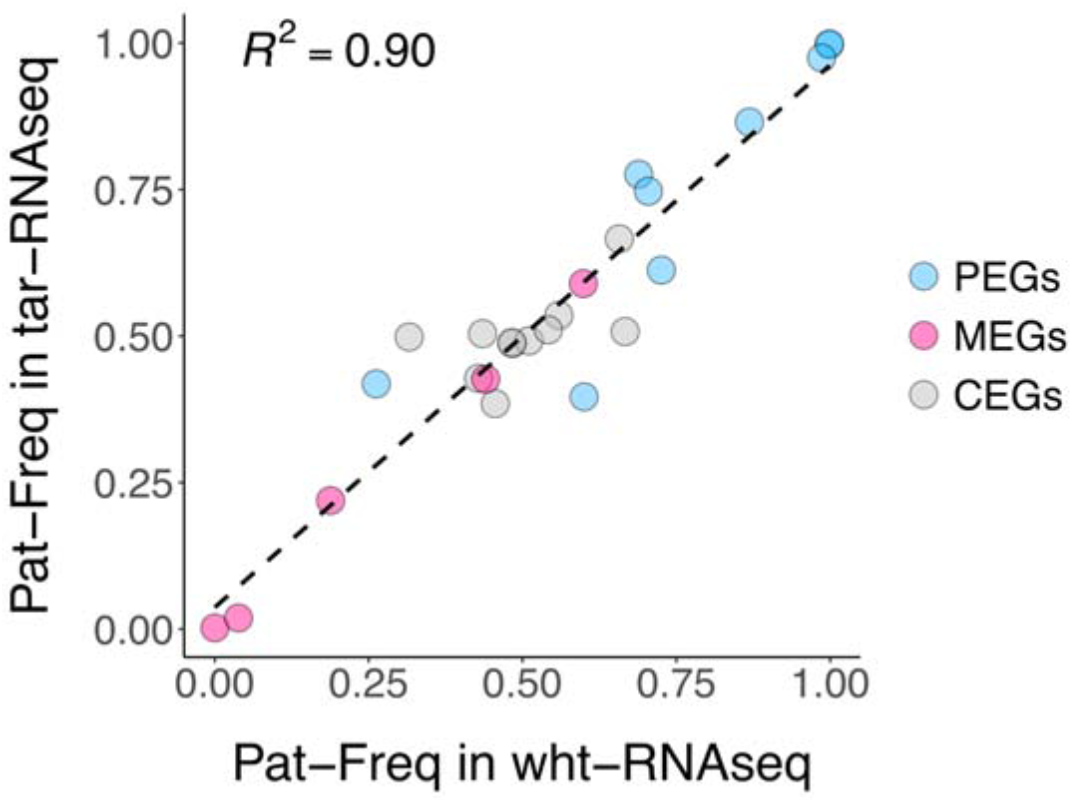
Correlation of gene-level average Pat-Freq between wht-RNAseq and tar-RNAseq. The scatter plot shows the Pat-Freq in each gene that had data in a sample sequenced with both wht-RNAseq and tar-RNAseq. PEGs, MEGs and CEGs are denoted by blue, pink and gray colors, respectively. The linear regression is shown by a dashed line.

RNA preparations from the same samples were processed with or without RiboErase to assess the reproducibility of ASE estimates with our workflows. The correlation of Ref-Freq between the samples in a pair was high (0.96, p < 2×10^−303^, n=2) (Supplementary Figure 4). Thus, similar to the well documented high reproducibility of whole Exome targeting^44^, tar-RNAseq delivers repeatable ASE estimates and tolerates the inclusion of RiboErase. Furthermore, validation of a subset of 19 SNPs in 3 samples revealed high correlation of Ref-Freq between ASE determined by pyrosequencing and tar-RNAseq (r=0.98, p < 1×10^−18^), similar to what is observed using wht-RNAseq in high RIN samples in mice (0.91, p < 1×10^−16^)^45^.

### Targeted RNA-seq improved SNP coverage

Even though ASE measurements between the two datasets were congruent, we obtained substantial improvement in SNP coverage from tar-RNAseq compared to wht-RNAseq. After removing the SNPs that had fewer than 10 total read counts, wht-RNAseq was able to cover only 79 (or 20%) of the SNPs, on average, across the samples, while tar-RNAseq covered up to 337 (or 80%) of the SNPs (Figure 3A) in the 75 genes that were common between the two datasets. We observed the same pattern when analyzing all SNPs in each dataset (Supplementary Figure 1). The improvement in coverage was achieved for tar-RNAseq with a mean of only 80 × 10^6^ reads per sample (SD 52 × 10^6^)--far less than the mean number of reads per sample of 269 × 10^6^ for wht-RNAseq (SD 110 × 10^6^). Stated in terms of number of total SNPs covered per billion bases sequenced, tar-RNAseq produced 54 SNPs/10^9^ bases while wht-RNAseq delivered only 2 SNPs/10^9^ bases. This improved use of sequencing resources was expected from the enrichment of the RNA fragments of interest. In our tar-RNAseq dataset, the percentage of bases that aligned to the targeted region was 85%, on average, indicative of successful enrichment (Methods). In principle, increasing sequencing depth could overcome the coverage deficits of wht-RNAseq. We calculate that the per sample cost would increase 35-fold, which for most projects is prohibitive especially in the context of a population study.

**Figure 3.**
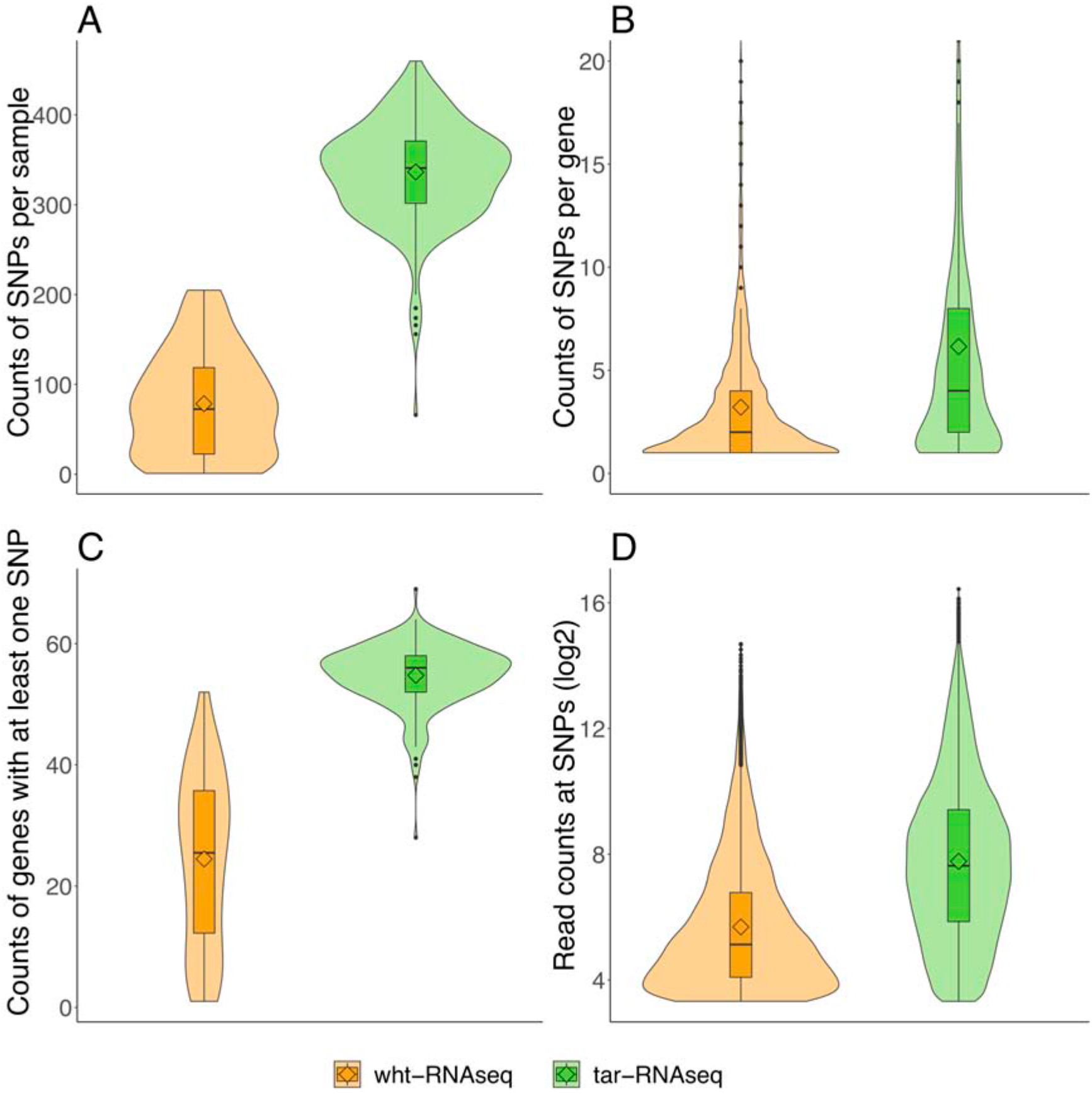
Comparison of SNP coverage between wht-RNAseq and tar-RNAseq. The violin plots show the distributions of (a) the counts of SNPs per sample, (b) the counts of SNPs per gene, (c) the counts of genes with at least one SNP, and (d) the read counts at SNPs. Orange and green colors denote wht-RNAseq and tar-RNAseq, respectively.

**Figure 4.**
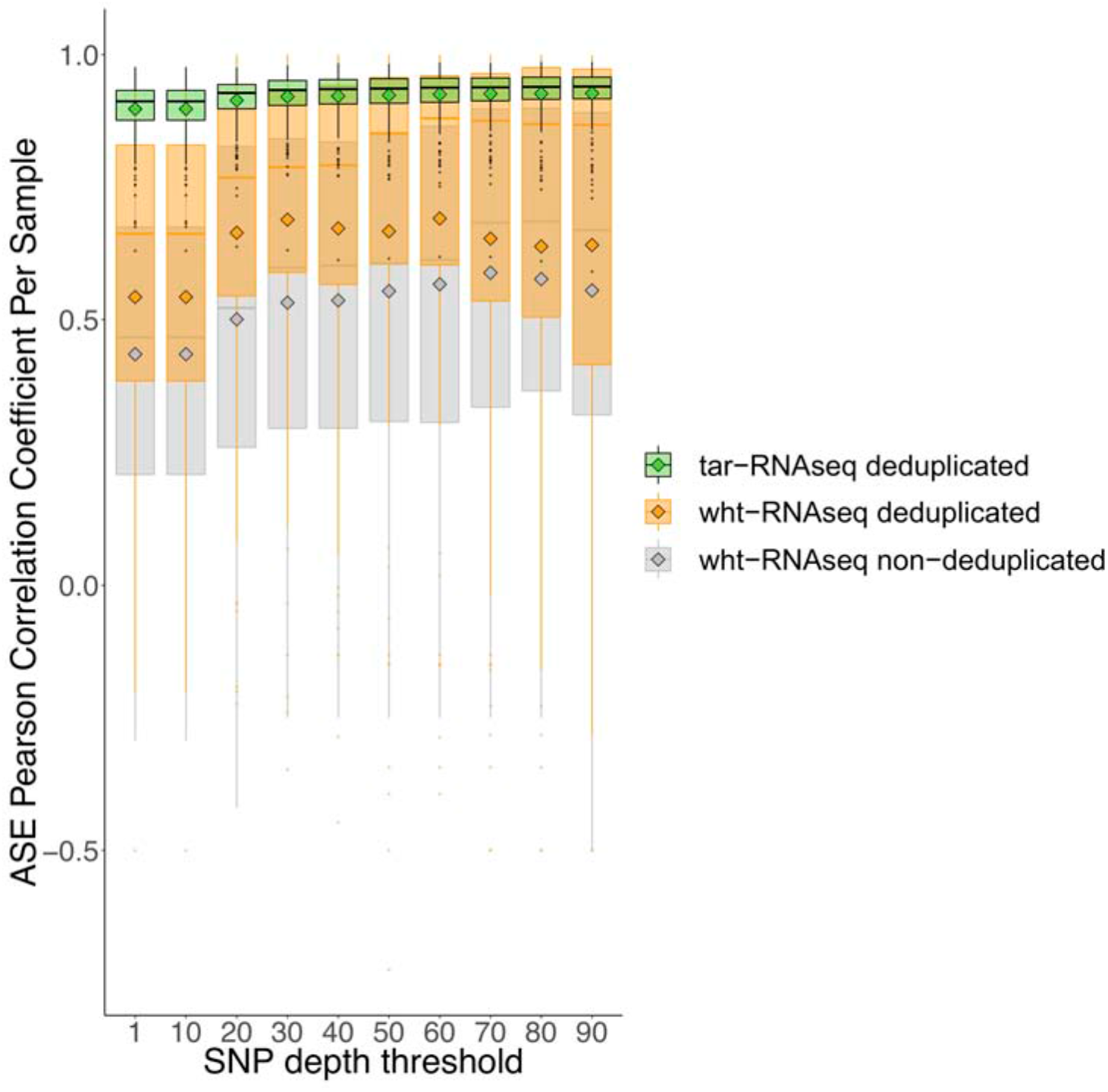
Concordance of ASE for SNPs in the same gene. The Pearson correlation coefficients were calculated from pairwise combinations of the SNPs from the same genes and their distributions are shown in box plots, stratified by escalated depth filtering thresholds. The mean of each distribution is indicated by a diamond. Data from tar-RNAseq (with deduping), wht-RNAseq (with deduping), and wht-RNAseq (without deduping) are denoted by green, orange and gray colors, respectively.

While both datasets had at least one SNP in the 75 genes analyzed here, each gene had informative SNPs in more samples in tar-RNAseq than in wht-RNAseq. Specifically, after quality filtration, 24 genes per sample had at least one SNP in wht-RNAseq with each gene carrying, on average, 3 SNPs; these values increased to 55 genes with 6 SNPs per gene in tar-RNAseq (Figure 3B, C). In addition, the average total read count for the final SNPs in tar-RNAseq was four times higher than in wht-RNAseq (Figure 3D). In sum, more informative SNPs were obtained by tar-RNAseq than wht-RNAseq, which allowed us to measure the inter-individual variability of ASE in more genes.

Arguably, the comparison could be confounded by the fact that different samples were analyzed between our tar-RNAseq and wht-RNAseq datasets as only one individual was sequenced by both technologies. However, it is unlikely that the difference in the samples between the two datasets was responsible for the large difference in the SNP coverage described above. Three sample-level QC metrics that could contribute to SNP coverage are number of genotyped SNPs, RIN, and maternal contamination. We show that RIN and maternal contamination did not differ significantly between the two datasets (Supplementary Table 2). We show that the choice of sequencing approach is more important than the number of genotyped SNPs through linear regressions in which the dependent variables were five different measures relevant to SNP coverage and the independent variables were tar-RNAseq (versus wht-RNAseq) and counts of genotyped exonic hetSNPs per sample (Supplementary Figure 3). Tar-RNAseq yielded a huge improvement in SNP coverage relative to wht-RNAseq at all observed numbers of genotyped SNPs. Moreover, even in the samples that had about 500 hetSNPs using wht-RNAseq, the SNP coverage was lower than in the samples that had about 300 hetSNPs using tar-RNAseq (Supplementary Figure 3). The mean number of genotyped SNPs was 418 for tar-RNAseq and 386 for wht-RNAseq, but evidently this difference could not underlie the improvement in SNP coverage using tar-RNAseq. In sum, our findings are not sensitive to the difference in the samples used in the two sequencing approaches.

### Targeted RNA-seq improved concordance of ASE from the same genes

Low SNP read coverage can limit the concordance of ASE between the SNPs from the same gene^27^. To determine the relationship between read coverage and concordance of ASE in our data, we calculated Pearson correlation coefficients for the pairwise combinations of SNPs mapping to the same gene in each sample. The mean correlation coefficient was 0.54 for wht-RNAseq and 0.90 for tar-RNAseq. At every read depth threshold, including the highest, the concordance was always much stronger in tar-RNAseq than in wht-RNAseq data (Figure 4). The concordance of the SNPs from the same exon showed the same pattern (Supplementary Figure 2). In wht-RNAseq, deduping improved SNP concordance but not to the level observed with deduped tar-RNAseq. Thus, hybridization capture improved this variable well beyond what would be expected from the increase in sequencing depth alone.

### Targeted RNA-seq in combination with rRNA depletion permitted assessment of ASE even in degraded samples

RNA degradation contributed to the reduced SNP coverage in the wht-RNAseq samples and inefficient rRNA removal is a factor known to interfere with the complexity of sequencing libraries^46^. In tar-RNAseq, ribosomal RNA should, in principle, have been removed by the hybridization reaction. However, we were able to rescue samples having low RIN by using rRNA depletion to improve SNP coverage in tar-RNAseq. Thus, we observed a strong positive correlation between RIN and SNP coverage fraction in the wht-RNAseq but not in the tar-RNAseq data (Figure 5). The combination of tar-RNAseq and rRNA depletion routinely produced high SNP coverage in samples with substantial RNA degradation (DV_200_ ~50%).

**Figure 5.**
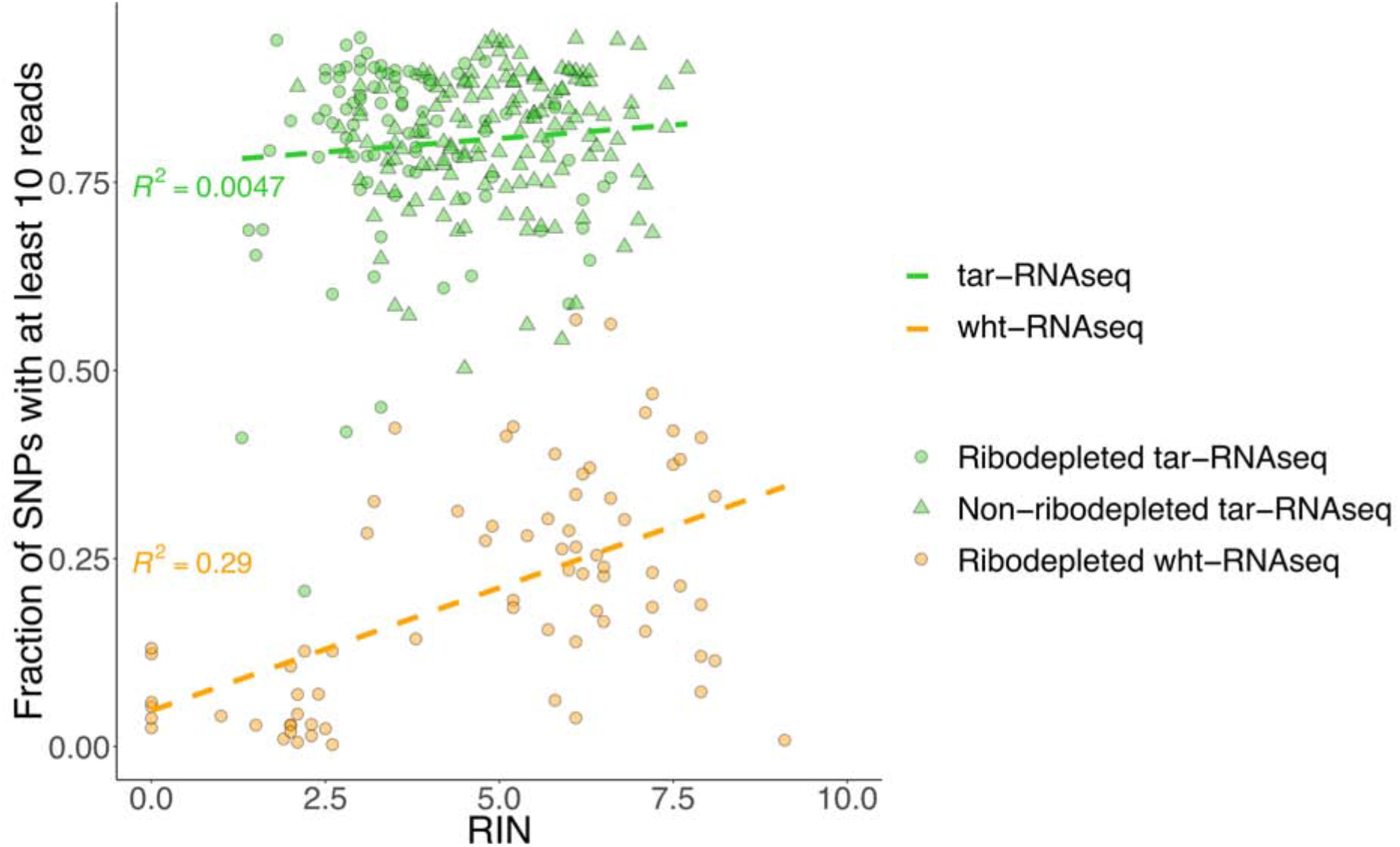
Combination of tar-RNAseq and RiboErase rescued most low RIN samples. The scatter plots show the fraction of SNPs covered by no fewer than 10 reads versus RIN. The ribodepleted tar-RNAseq, non-ribodepleted tar-RNAseq, and ribodepleted wht-RNAseq samples are denoted by green circles, green triangles, and orange circles, respectively. The linear regression for tar-RNAseq and wht-RNAseq data points is shown by a green and an orange dashed line, respectively.

### Targeted RNA-seq improved the measurement of relative expression

Although the focus of our efforts was on ASE and not relative expression^45^, we compared the performance of the two approaches with particular regard to degraded samples. As expected, compared with wht-RNAseq, tar-RNAseq yielded higher TPM values as the genes of interest constituted a larger fraction of the total sequenced transcripts (Supplementary Figure 5). Expression measurements were more consistent for low RIN samples from tar-RNAseq than for wht-RNAseq (Supplementary Figures 5A and 6B). Below a RIN of 3, many genes were expressed at TPM close to 0 in the wht-RNAseq data set--visible as a transition in the heat maps of gene expression versus RIN (Supplementary figures 5A and 5B). No such transition was seen for the tar-RNAseq heat map and the relative expression of highly or lowly expressed genes was not influenced by RNA integrity. These observations indicate that tar-RNAseq produced more reliable measurements of expression and was less affected by sample quality.

## Discussion

ASE analysis has been performed on a variety of wht-RNAseq datasets including simulated sequences^47^, RNA from cells cultured in vitro^48,49^, and RNA from inbred mice^45^. In humans, ASE analysis of RNA from many tissues was performed as part of the GTEx project^4,5^ (https://gtexportal.org/home/). Placentas were not included in GTEx, but human placental tissue has been analyzed by other groups using wht-RNAseq and analyzed for ASE^18,19,50^. The goal of these studies in regard to ASE was to identify imprinted genes through a transcriptome wide approach and to categorize them by imprinting status. The ENCODE study aimed to go beyond categorization and pursued a more quantitative approach that entailed calculation of the significance values for the parent of origin effect for individual SNPs^14^. In contrast with the foregoing studies, our goal was to generate quantitative gene-level ASE estimates with high precision and accuracy. Such estimates are required for investigation of the functional significance of inter-individual variation in ASE^29^.

Toward that end, we compared allelic count distributions in tar-RNAseq and wht-RNAseq datasets for a set of 75 genes that had been reported in the literature to be imprinted and that had at least one SNP in both datasets. We found that the two methods produced similar allelic expression biases. However, wht-RNAseq was able to cover only 20% of the SNPs, on average, across the samples, whereas tar-RNAseq covered 80% of the SNPs, with improvements in SNP coverage of 27-fold per billion bases sequenced. In humans, the paucity of SNPs makes it imperative to cover all the SNPs in the genes of interest. Using tar-RNAseq, we were able to obtain sufficient coverage at four times as many SNPs in twice as many genes in a sample, on average. Moreover, the mean number of SNPs per gene doubled, and the mean read depth per SNP increased four-fold, without increasing library preparation costs, making tar-RNAseq more cost effective (Supplementary Table 3). A complete data set would have sufficient reads at every SNP in every person, which is a goal that was more closely achieved by tar-RNA seq than by wht-RNAseq. Having a richer dataset will enable us to determine the interindividual variation in ASE for more genes across more individuals, so that we can better query the association between genomic imprinting and growth phenotypes in our cohort study.

Maternal contamination is a potential confounder unique to placental tissue and is a limiting factor in molecular analyses^51^. The degree of contamination can be directly determined from the RNAseq data by quantitating non-fetal alleles^19^. The greater SNP sequencing coverage and depth of SNPs in tar-RNAseq enabled us to quantify maternal contamination for each gene in each placenta with greater sensitivity.

Importantly, targeted RNA-seq had some additional non-anticipated benefits. Gene-level ASE estimates are imprecise due to poor concordance of SNP-level ASE over a gene. Some discordance between the SNPs can be due to differing imprinting status between the transcript variants from the same gene^45^, but poorly identified technical variables also contribute^4,5^. It is known that selecting SNPs with higher sequencing coverage leads to improved concordance^27^, which we also saw in wht-RNAseq. Removing PCR duplicates further improved concordance but not nearly to the levels achieved by tar-RNAseq. In contrast with relative expression analyses, ASE is only based on the read count ratio between the alleles and removing read duplicates reduces the technical noise. The greatly improved concordance of SNPs strongly argues in favor of using tar-RNAseq for applications that require accurate gene-level ASE estimates. In future efforts, it might be possible to gain additional power for SNPs with low read depth by using unique molecular indices (UMI)^52^ in conjunction with tar-RNAseq as there are reports that PCR duplicates can affect ASE quantification in such circumstances^21^.

Degraded RNA is found in many human samples, including term placentas, post-mortem samples of stomach and kidney, and formalin-fixed paraffin-embedded (FFPE) samples^51,53,54^ (https://gtexportal.org/home/). We were successful in combining random primed library preparation with rRNA depletion to generate libraries from degraded RNA for tar-RNAseq. Little to no loss of coverage was observed with degraded RNA from most samples, and we salvaged samples with DV_200_ as low as 50%. The critical role for efficient rRNA depletion in preparing libraries from degraded RNA is well known^46^ and is in part due to the inability to target the poly(A) tail in degraded samples. The hybridization reaction with the capturing oligos should, in principle, be sufficient to remove rRNA. However, our results show that in degraded samples, removal of rRNA prior to library preparation improved SNP coverage. Improved data quality has previously been reported for gene expression analysis in FFPE samples with RNA-seq and rRNA depletion^55^ or when capturing the whole exome^56^. However, to our knowledge, our study is the first to report the unexpected synergy between rRNA depletion and tar-RNAseq. It was also more efficient and cost effective to focus on a targeted region of only 3 Mb instead of the human exome of 64 Mb (the total length of Roche SeqCap EZ Exome Probes). Tar-RNAseq also improved relative expression estimates for degraded samples. Importantly, we document that cost savings is only one of the advantages of tar-RNAseq and other synergies may become the predominant motive to use this technology as sequencing costs continue to fall.

A strength of our study is that the placenta samples were collected from healthy women who were of similar ages and belonged to the same cohort and ethnicity, using a standardized protocol. Moreover, we compared the same 75 genes using both wht-RNAseq and tar-RNAseq. A limitation of our study is that only one sample was sequenced using both methods. However, we examined three parameters that could potentially differ between samples and influence data yield (RIN, maternal contamination, number of genotyped SNPs) and showed that our conclusion about the superiority of tar-RNAseq was not sensitive to any of these parameters. We also note that library preparation reagents for wht-RNAseq libraries and tar-RNAseq libraries were from different manufacturers. Although we did not try to estimate ASE in relation to the cellular composition of the fetal compartment of the placenta, a recent single cell study showed that placental samples collected using a protocol similar to ours were mostly comprised of trophoblast and syncytiotrophoblast cells of the fetus^57^.

In conclusion, we compared tar-RNAseq and wht-RNAseq in a study of ASE in 75 known imprinted genes in placental tissue collected from a healthy human cohort. Tar-RNAseq covered more SNPs of interest and at greater depth. In previous research on humans, discordant ASE values for SNPs of the same gene have limited the ability to accurately quantify ASE. We show that Tar-RNAseq improved the reliability of ASE detection by greatly increasing the concordance of ASE measurement between the SNPs from the same gene. In combination with rRNA depletion, tar-RNAseq performed well even in cases of degraded RNA. The advantages of tar-RNAseq go beyond the savings on sequencing costs alone and include higher accuracy in ASE estimates in samples with varying RNA quality, as is typical for field collections. Targeted sequencing will benefit the study of associations between individual variation in ASE and phenotypes in humans or in other species where growth phenotypes are of interest, such as domesticated animals. The data we presented here originated from field samples and provide metrics to inform the design of such projects.

## Supporting information

Supplementary .bed file

## Supplementary information

The supplementary materials were deposited to Figshare (DOI: 10.6084/m9.figshare.13637843).

## Acknowledgments

This research was supported by the Eunice Kennedy Shriver National Institute of Child Health & Human Development of the National Institutes of Health (R01HD088521 and R21HD077465 to BIS); the John Templeton Foundation (52269 to BIS); and the National Science Foundation program in Biological Anthropology (NSF BCS-1354814 to BIS). The content is solely the responsibility of the authors and does not necessarily represent the official views of the National Institutes of Health, the John Templeton Foundation, or the National Science Foundation. The authors thank the study participants who made this research possible as well as three Malian gynecologists for their advice and helpful suggestions: Prof. Amadou Dolo, Prof. Niani Mounkoro, and Prof. Mamadou Traoré. We also thank our field manager, Zachary Dolo, and the Malian medical team: Dr. Gouro Dicko, Dr. Akoro Dolo, Madeleine Goita, Aissa Dolo, Younus Dolo, Geremy Sagara, and Safoura Guindo. For permission to carry out this study in Mali, we are grateful to the Centre National de la Recherche Scientifique et Technologique and the Comité d’Ethique de la Faculté de Médecine de Pharmacie et d’Onto-Stomatologie of the University of Sciences, Techniques, and Technologies of Bamako (authorization N°2016/68/CD/FMPOS). We also acknowledge the contribution of the Advanced Genomics Core at the University of Michigan.

## Data access

The data for the wht-RNAseq study are registered in dbGap as “Placental Transcriptome and Stunting.” The wht-RNAseq data and the corresponding genotypes obtained through targeted sequencing, the FASTQ files with the sequences from RNAseq, and the SNP level file with the allele specific counts were deposited in dbGaP as phs001782.v1.p1. The person level non-molecular data is available at the same site. The analogous data for the tar-RNAseq will be made available through dbGaP once the manuscript has been accepted.

## References

1. Tucci V, Isles AR, Kelsey G, et al. Genomic Imprinting and Physiological Processes in Mammals. Cell. 2019;176(5):952–965. doi:10.1016/j.cell.2019.01.043

2. Batista RA, Köhler C. Genomic imprinting in plants-revisiting existing models. Published online 2020. doi:10.1101/gad.332924

3. Moore T, Haig D. Genomic imprinting in mammalian development: a parental tug-of-war. Trends Genet. 1991;7(2):45–49. doi:10.1016/0168-9525(91)90230-N

4. Baran Y, Subramaniam M, Biton A, et al. The landscape of genomic imprinting across diverse adult human tissues. Genome Res. 2015;25(7):927–936. doi:10.1101/gr.192278.115

5. Babak T, Deveale B, Tsang EK, et al. Genetic conflict reflected in tissue-specific maps of genomic imprinting in human and mouse. Nat Genet. 2015;47(5):544–549. doi:10.1038/ng.3274

6. Peters J. The role of genomic imprinting in biology and disease: An expanding view. Nat Rev Genet. 2014;15(8):517–530. doi:10.1038/nrg3766

7. Farhadova S, Gomez-Velazquez M, Feil R. Stability and lability of parental methylation imprints in development and disease. Genes (Basel). 2019;10(12). doi:10.3390/genes10120999

8. Thamban T, Agarwaal V, Khosla S. Role of genomic imprinting in mammalian development. J Biosci. 2020;45(1). doi:10.1007/s12038-019-9984-1

9. Li Y, Li J. Technical advances contribute to the study of genomic imprinting. PLoS Genet. 2019;15(6). doi:10.1371/journal.pgen.1008151

10. Wang X, Clark AG. Using next-generation RNA sequencing to identify imprinted genes. Heredity (Edinb). 2014;113(2):156–166. doi:10.1038/hdy.2014.18

11. Wang X, Miller DC, Harman R, Antczak DF, Clark AG. Paternally expressed genes predominate in the placenta. Proc Natl Acad Sci U S A. 2013;110(26):10705–10710. doi:10.1073/pnas.1308998110

12. Wang Y, Gao S, Zhao Y, et al. Allele-specific expression and alternative splicing in horse×donkey and cattle×yak hybrids. Zool Res. 2019;40(4):293–304. doi:10.24272/j.issn.2095-8137.2019.042

13. Chen Z, Hagen DE, Wang J, et al. Global assessment of imprinted gene expression in the bovine conceptus by next generation sequencing. Epigenetics. 2016;11(7):501–516. doi:10.1080/15592294.2016.1184805

14. Zink F, Magnusdottir DN, Magnusson OT, et al. Insights into imprinting from parent-of-origin phased methylomes and transcriptomes. Nat Genet. 2018;50(11):1542–1552. doi:10.1038/s41588-018-0232-7

15. Metsalu T, Viltrop T, Tiirats A, et al. Using RNA sequencing for identifying gene imprinting and random monoallelic expression in human placenta. Epigenetics. 2014;9(10):1397–1409. doi:10.4161/15592294.2014.970052

16. Jadhav B, Monajemi R, Gagalova KK, et al. RNA-Seq in 296 phased trios provides a high-resolution map of genomic imprinting. BMC Biol. 2019;17(1). doi:10.1186/s12915-019-0674-0

17. Mozaffari S V., Stein MM, Magnaye KM, Nicolae DL, Ober C. Parent of origin gene expression in a founder population identifies two new candidate imprinted genes at known imprinted regions. PLoS One. 2018;13(9):e0203906. doi:10.1371/journal.pone.0203906

18. Pilvar D, Reiman M, Pilvar A, Laan M. Parent-of-origin-specific allelic expression in the human placenta is limited to established imprinted loci and it is stably maintained across pregnancy. Clin Epigenetics. 2019;11(1):94. doi:10.1186/s13148-019-0692-3

19. Hamada H, Okae H, Toh H, et al. Allele-Specific Methylome and Transcriptome Analysis Reveals Widespread Imprinting in the Human Placenta. Am J Hum Genet. 2016;99(5):1045–1058. doi:10.1016/j.ajhg.2016.08.021

20. Gulyás-Kovács A, Keydar I, Xia E, et al. Unperturbed expression bias of imprinted genes in schizophrenia. Nat Commun. 2018;9(1):7. doi:10.1038/s41467-018-04960-9

21. Castel SE, Levy-Moonshine A, Mohammadi P, Banks E, Lappalainen T. Tools and best practices for data processing in allelic expression analysis. Genome Biol. 2015;16(1):195. doi:10.1186/s13059-015-0762-6

22. Stevenson KR, Coolon JD, Wittkopp PJ. Sources of bias in measures of allele-specific expression derived from RNA-seq data aligned to a single reference genome. BMC Genomics. 2013;14(1):536. doi:10.1186/1471-2164-14-536

23. Van De Geijn B, Mcvicker G, Gilad Y, Pritchard JK. WASP: Allele-specific software for robust molecular quantitative trait locus discovery. Nat Methods. 2015;12(11):1061–1063. doi:10.1038/nmeth.3582

24. Nothnagel M, Wolf A, Herrmann A, et al. Statistical inference of allelic imbalance from transcriptome data. Hum Mutat. 2011;32(1):98–106. doi:10.1002/humu.21396

25. Heap GA, Yang JHM, Downes K, et al. Genome-wide analysis of allelic expression imbalance in human primary cells by high-throughput transcriptome resequencing. Hum Mol Genet. 2009;19(1):122–134. doi:10.1093/hmg/ddp473

26. Fontanillas P, Landry CR, Wittkopp PJ, et al. Key considerations for measuring allelic expression on a genomic scale using high-throughput sequencing. Mol Ecol. 2010;19(SUPPL. 1):212–227. doi:10.1111/j.1365-294X.2010.04472.x

27. Zou J, Hormozdiari F, Jew B, et al. Leveraging allelic imbalance to refine fine-mapping for eQTL studies. PLoS Genet. 2019;15(12):e1008481. doi:10.1371/journal.pgen.1008481

28. DeVeale B, van der Kooy D, Babak T. Critical evaluation of imprinted gene expression by RNA-seq: A new perspective. PLoS Genet. 2012;8(3). doi:10.1371/journal.pgen.1002600

29. Vincenz C, Lovett JL, Wu W, Shedden K, Strassmann BI. Loss of Imprinting in Human Placentas Is Widespread, Coordinated, and Predicts Birth Phenotypes. Mol Biol Evol. 2020;37(2). doi:10.1093/molbev/msz226

30. Bolger AM, Lohse M, Usadel B. Trimmomatic: A flexible trimmer for Illumina sequence data. Bioinformatics. 2014;30(15):2114–2120. doi:10.1093/bioinformatics/btu170

31. Li H, Durbin R. Fast and accurate short read alignment with Burrows-Wheeler transform. Bioinformatics. 2009;25(14):1754–1760. doi:10.1093/bioinformatics/btp324

32. Depristo MA, Banks E, Poplin R, et al. A framework for variation discovery and genotyping using next-generation DNA sequencing data. Nat Genet. 2011;43(5):491–501. doi:10.1038/ng.806

33. Van der Auwera GA, Carneiro MO, Hartl C, et al. From fastQ data to high-confidence variant calls: The genome analysis toolkit best practices pipeline. Curr Protoc Bioinforma. 2013;43(SUPL.43). doi:10.1002/0471250953.bi1110s43

34. Karimzadeh M, Ernst C, Kundaje A, Hoffman MM. Umap and Bismap: Quantifying genome and methylome mappability. Nucleic Acids Res. 2018;46(20). doi:10.1093/nar/gky677

35. Amemiya HM, Kundaje A, Boyle AP. The ENCODE Blacklist: Identification of Problematic Regions of the Genome. Sci Rep. 2019;9(1). doi:10.1038/s41598-019-45839-z

36. Bailey JA, Gu Z, Clark RA, et al. Recent segmental duplications in the human genome. Science (80-). 2002;297(5583):1003–1007. doi:10.1126/science.1072047

37. Panousis NI, Gutierrez-Arcelus M, Dermitzakis ET, Lappalainen T. Allelic mapping bias in RNA-sequencing is not a major confounder in eQTL studies. Genome Biol. 2014;15(9):467. doi:10.1186/s13059-014-0467-2

38. Kim D, Langmead B, Salzberg SL. HISAT: A fast spliced aligner with low memory requirements. Nat Methods. 2015;12(4):357–360. doi:10.1038/nmeth.3317

39. Harrow J, Frankish A, Gonzalez JM, et al. GENCODE: The reference human genome annotation for the ENCODE project. Genome Res. 2012;22(9):1760–1774. doi:10.1101/gr.135350.111

40. Van De Geijn B, Mcvicker G, Gilad Y, Pritchard JK. WASP: Allele-specific software for robust molecular quantitative trait locus discovery. Nat Methods. 2015;12(11):1061–1063. doi:10.1038/nmeth.3582

41. Pertea M, Kim D, Pertea GM, Leek JT, Salzberg SL. Transcript-level expression analysis of RNA-seq experiments with HISAT, StringTie and Ballgown. Nat Protoc. 2016;11(9):1650–1667. doi:10.1038/nprot.2016.095

42. Majewska M, Lipka A, Paukszto L, et al. Transcriptome profile of the human placenta. Funct Integr Genomics. 2017;17(5):551–563. doi:10.1007/s10142-017-0555-y

43. McLaren W, Gil L, Hunt SE, et al. The Ensembl Variant Effect Predictor. Genome Biol. 2016;17(1). doi:10.1186/s13059-016-0974-4

44. Cherukuri PF, Maduro V, Fuentes-Fajardo K V., et al. Replicate exome-sequencing in a multiple-generation family: improved interpretation of next-generation sequencing data. BMC Genomics. 2015;16(1):998. doi:10.1186/s12864-015-2107-y

45. Perez JD, Rubinstein ND, Fernandez DE, et al. Quantitative and functional interrogation of parent-of-origin allelic expression biases in the brain. Elife. 2015;4(JULY 2015):41. doi:10.7554/eLife.07860

46. Stark R, Grzelak M, Hadfield J. RNA sequencing: the teenage years. Nat Rev Genet. 2019;20(11):631–656. doi:10.1038/s41576-019-0150-2

47. Raghupathy N, Choi K, Vincent MJ, et al. Hierarchical analysis of RNA-seq reads improves the accuracy of allele-specific expression. In: Bioinformatics. Vol 34. Oxford University Press; 2018:2177–2184. doi:10.1093/bioinformatics/bty078

48. Gutierrez-Arcelus M, Lappalainen T, Montgomery SB, et al. Passive and active DNA methylation and the interplay with genetic variation in gene regulation. Elife. 2013;2013(2). doi:10.7554/eLife.00523

49. Lappalainen T, Sammeth M, Friedländer MR, et al. Transcriptome and genome sequencing uncovers functional variation in humans. Nature. 2013;501(7468):506–511. doi:10.1038/nature12531

50. Hanna CW, Peñaherrera MS, Saadeh H, et al. Pervasive polymorphic imprinted methylation in the human placenta. Genome Res. 2016;26(6):756–767. doi:10.1101/gr.196139.115

51. Konwar C, Del Gobbo G, Yuan V, Robinson WP. Considerations when processing and interpreting genomics data of the placenta. Placenta. 2019;84:57–62. doi:10.1016/j.placenta.2019.01.006

52. Islam S, Zeisel A, Joost S, et al. Quantitative single-cell RNA-seq with unique molecular identifiers. Nat Methods. 2014;11(2):163–166. doi:10.1038/nmeth.2772

53. Walker DG, Whetzel AM, Serrano G, Sue LI, Lue LF, Beach TG. Characterization of RNA isolated from eighteen different human tissues: results from a rapid human autopsy program. Cell Tissue Bank. 2016;17(3):361–375. doi:10.1007/s10561-016-9555-8

54. Zhang P, Lehmann BD, Shyr Y, Guo Y. The Utilization of Formalin Fixed-Paraffin-Embedded Specimens in High Throughput Genomic Studies. Int J Genomics. 2017;2017. doi:10.1155/2017/1926304

55. Zhao W, He X, Hoadley KA, Parker JS, Hayes DN, Perou CM. Comparison of RNA-Seq by poly (A) capture, ribosomal RNA depletion, and DNA microarray for expression profiling. BMC Genomics. 2014;15(1):419. doi:10.1186/1471-2164-15-419

56. Pennock ND, Jindal S, Horton W, et al. RNA-seq from archival FFPE breast cancer samples: Molecular pathway fidelity and novel discovery. BMC Med Genomics. 2019;12(1):195. doi:10.1186/s12920-019-0643-z

57. Yuan V, Hui D, Yin Y, Peñaherrera M, Beristain A, Robinson W. Cell-specic Characterization of the Placental Methylome. Published online September 12, 2020. doi:10.21203/rs.3.rs-38223/v2

