## Supplementary figures and images for "Targeted RNA-seq improves efficiency, resolution, and accuracy of allele specific expression for human term placentas"

### 6B

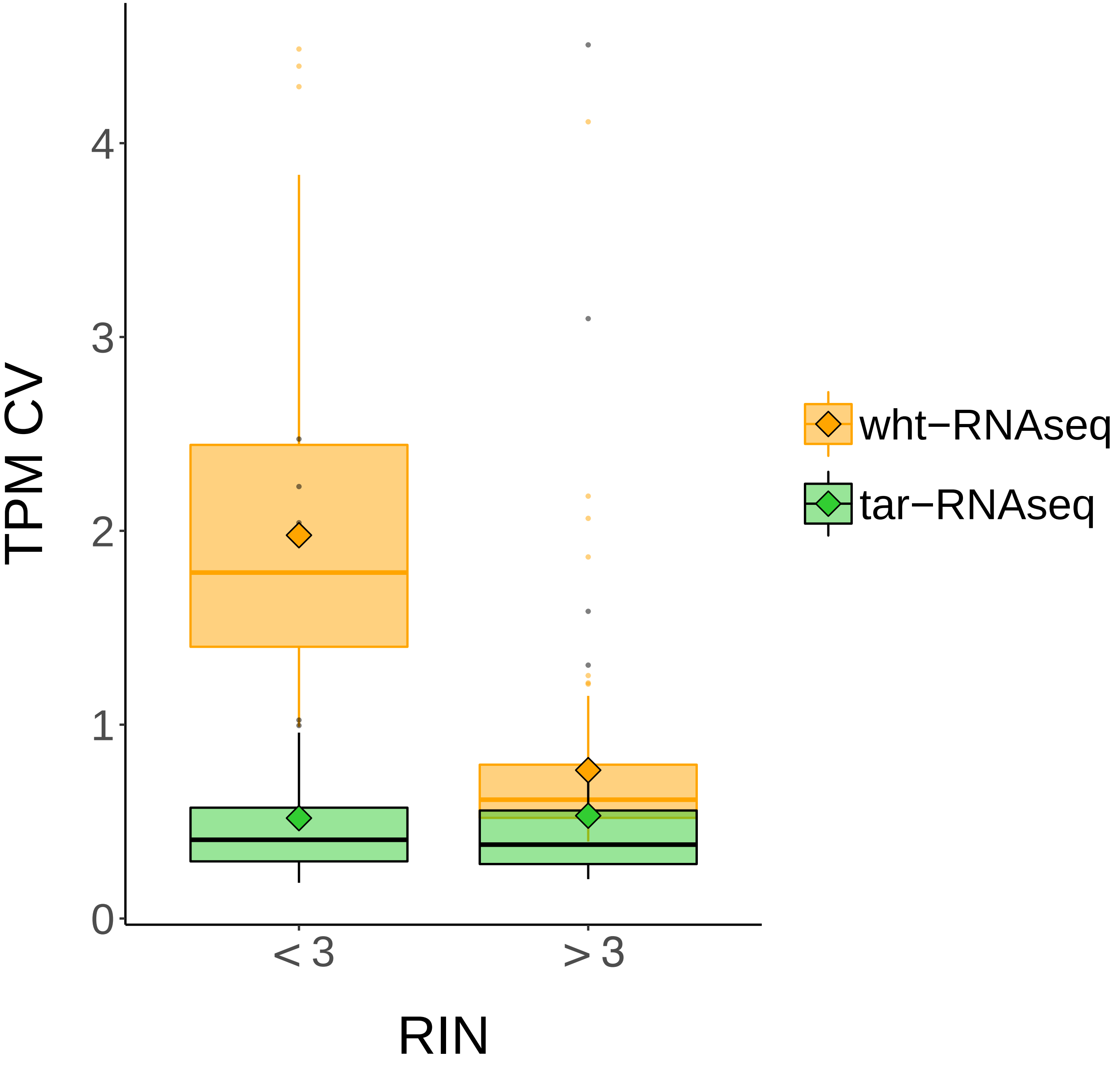

### Supplementary Figure 1

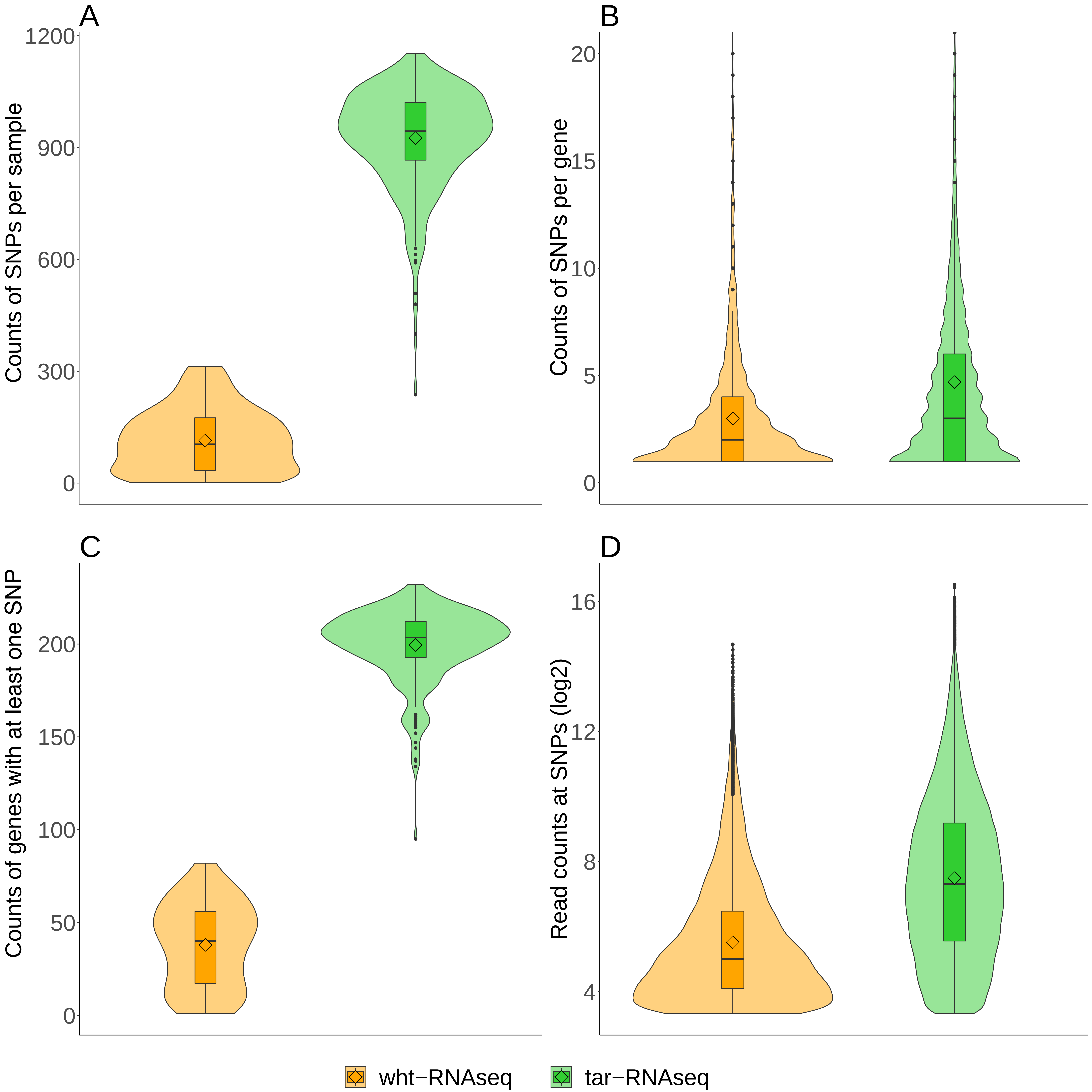

### Supplementary Figure 2

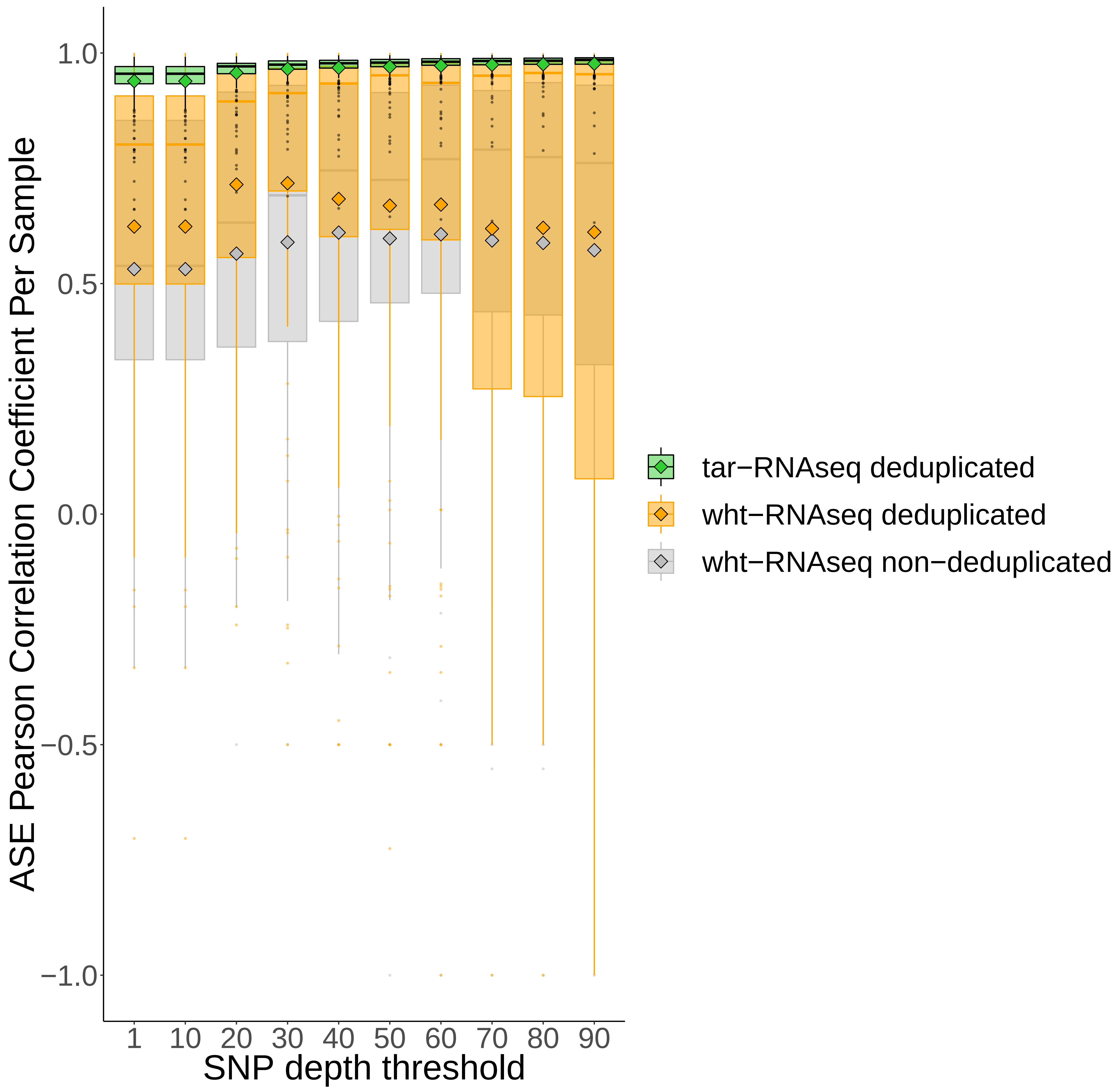

### Supplementary Figure 3

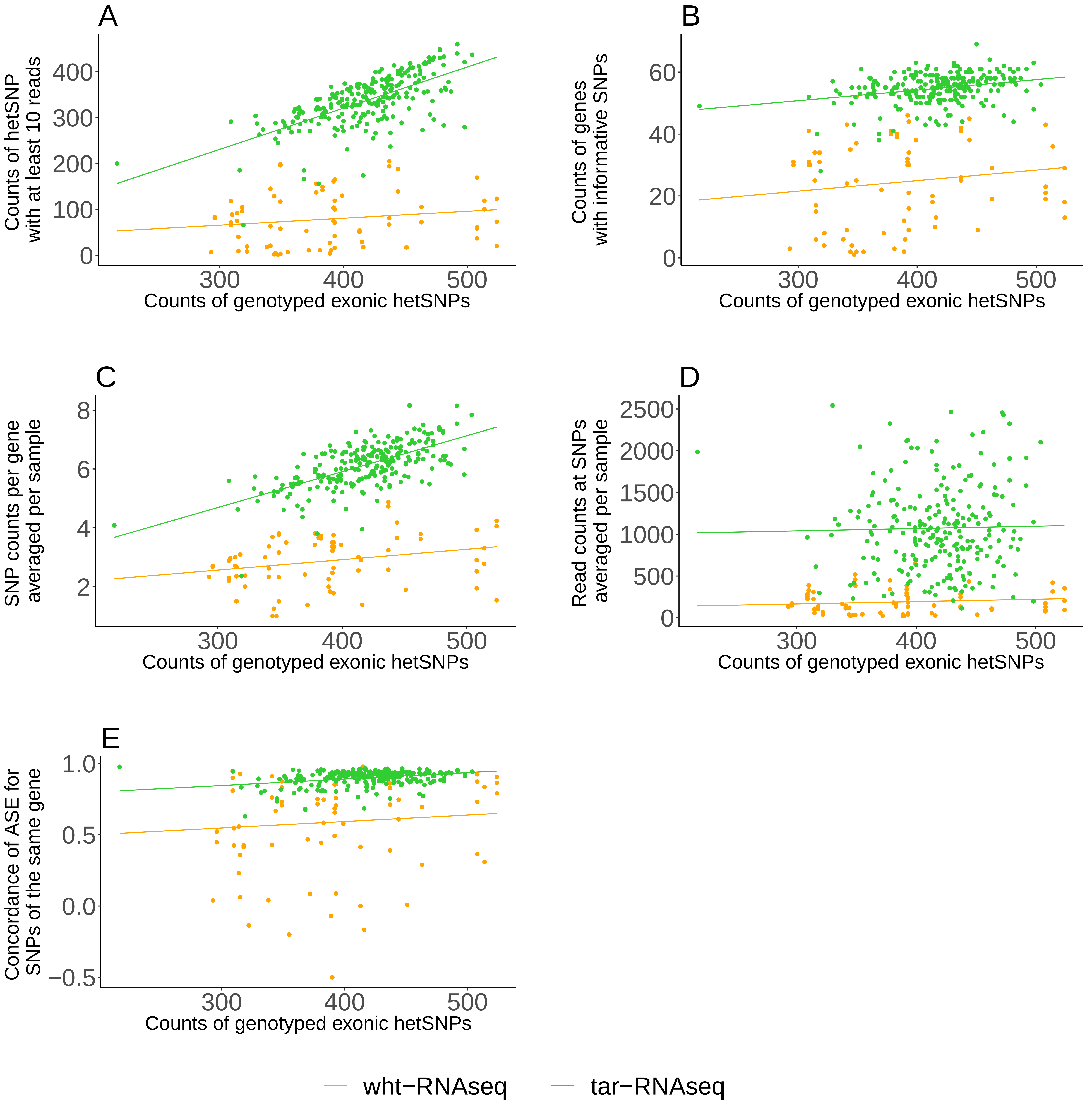

### Supplementary Figure 4

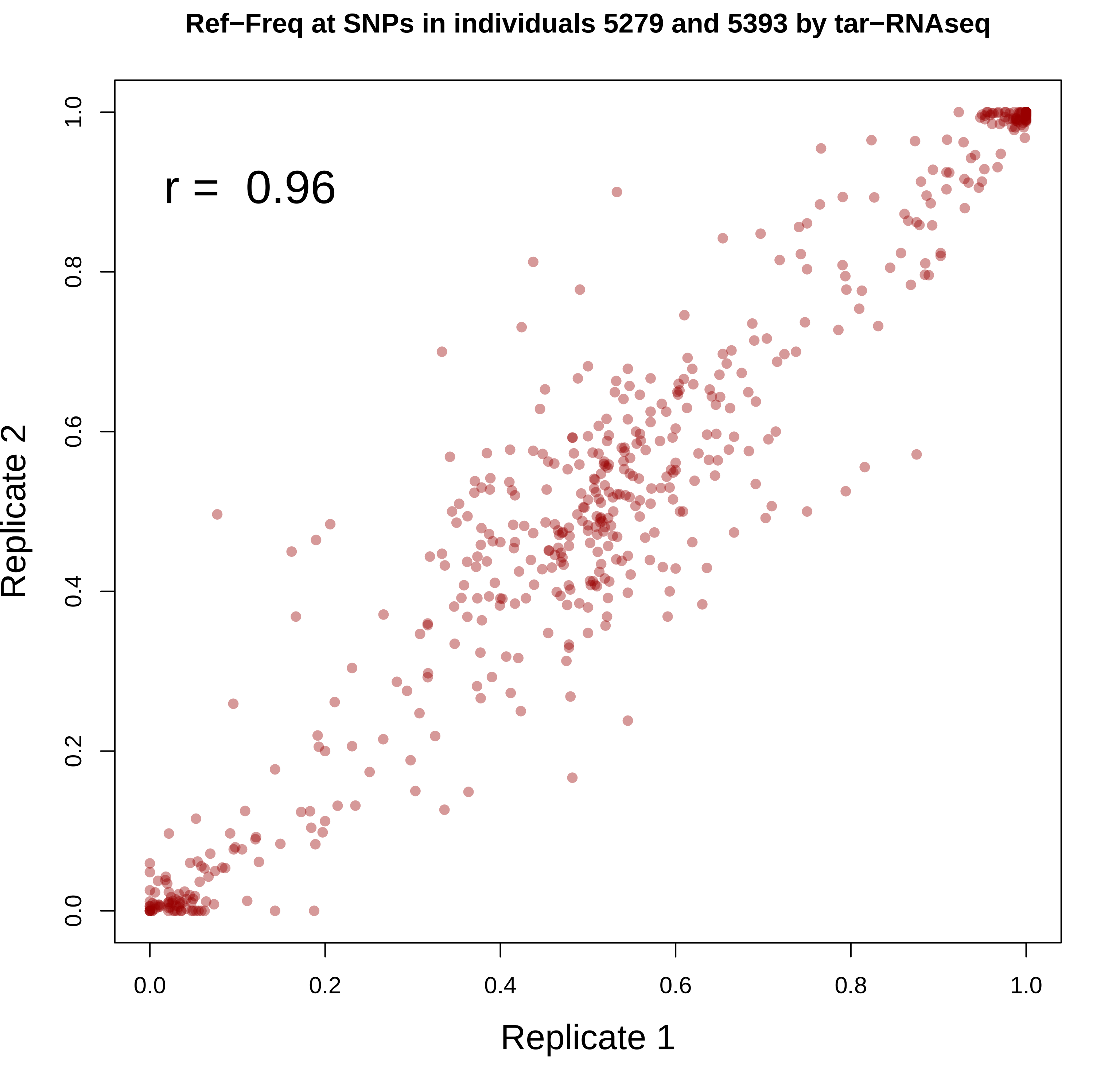

### Supplementary Figure 5

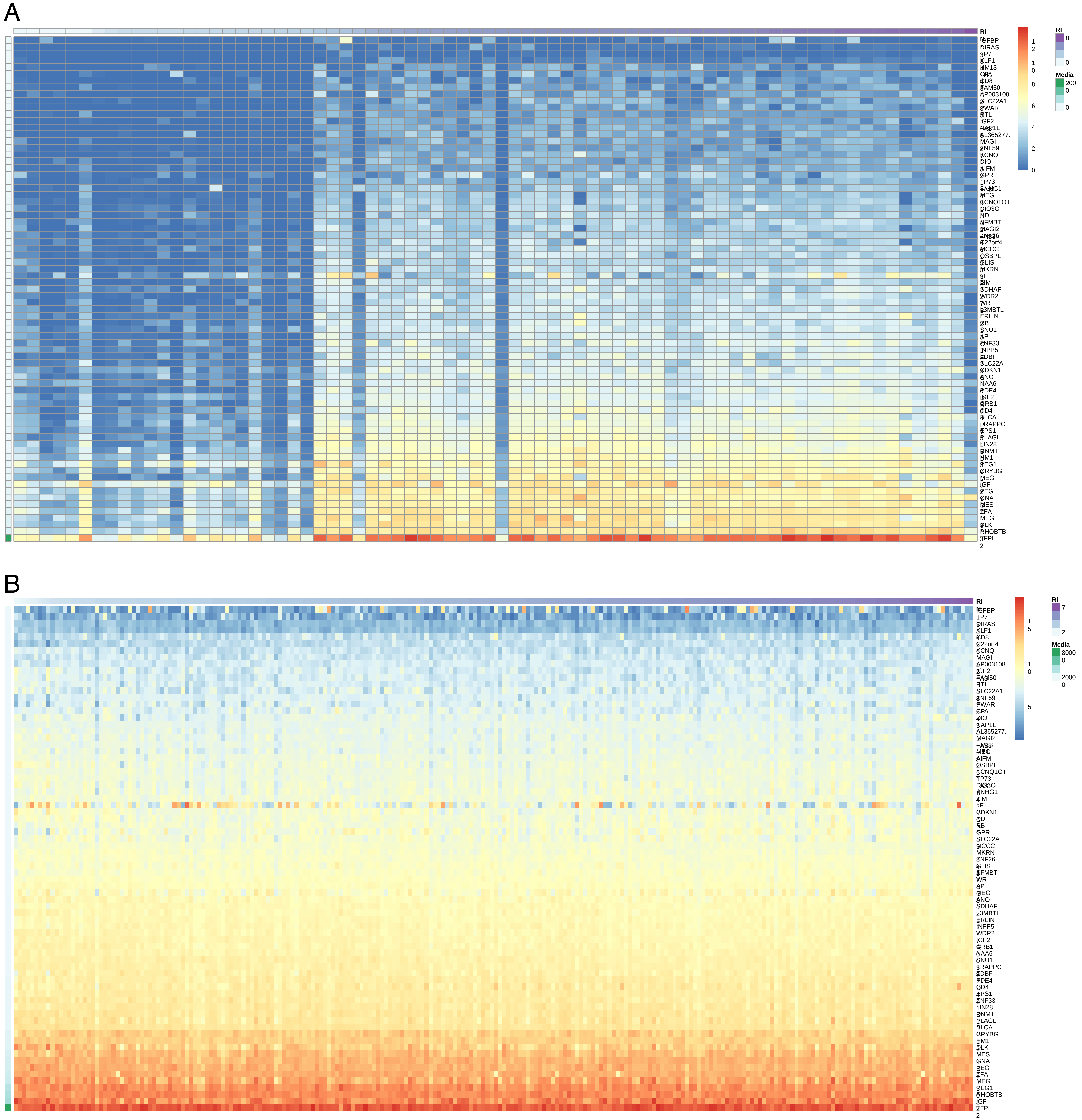
