## Supplementary material for "Targeted RNA-seq improves efficiency, resolution, and accuracy of allele specific expression for human term placentas": Supplentary Figure Legends

**Supplementary Figure 1.** Comparison of SNP coverage between wht-RNAseq and tar-RNAseq including all SNPs from genes in both data sets without regard to overlap or imprinting status. The violin plots show the distributions of (a) the counts of SNPs per sample, (b) the counts of SNPs per gene, (c) the counts of genes with at least one SNP, and (d) the read counts at SNPs. wht-RNAseq and tar-RNAseq are denoted by orange and green colors, respectively.

**Supplementary Figure 2.** Concordance of ASE for SNPs in the same exon. The Pearson correlation coefficients were calculated from pairwise combinations of the SNPs from the same exons and their distributions are shown in box plots, stratified by escalated depth filtering thresholds. The mean of each distribution is indicated by a diamond. Data from tar-RNAseq (with deduping), wht-RNAseq (with deduping), and wht-RNAseq (without deduping) are denoted by green, orange and gray colors, respectively.

**Supplementary Figure 3**. Improvement in SNP coverage and ASE concordance is driven by the targeting approach. Linear regression models were used to ascertain the effect of the targeting approach and the genotyping yield on the SNP coverage or the ASE concordance. The independent variables in each model are sequencing approach (tar-RNAseq or wht-RNAseq), and the number of genotyped exonic heterozygous SNPs for each sample. The dependent variables are: (a) the counts of SNPs per sample, (b) the counts of SNPs per gene, averaged for each sample, (c) the counts of genes with at least one SNP, averaged for each sample, (d) the read counts at SNPs, averaged for each sample, and (e) the Pearson correlation coefficients calculated from pairwise combinations of the SNPs from the same genes in each sample. In each plot, the dots are the observed data and the lines are the linear regression. Data from wht-RNAseq and tar-RNAseq are denoted by orange and green colors, respectively.

**Supplementary Figure 4.** Correlation of Ref-Freq for SNPs in biopsy replicates. The replicates were obtained from separate RNA isolates from biopsies of placenta ID 5279 and ID 5393 and the sequencing libraries in each pair were made with or without RiboErase.

**Supplementary Figure 5.** Relative gene expression measured using tar-RNAseq and wht-RNAseq. TPM values with log2 transformation (after adding 1) across 75 genes shown as heatmaps for (A) wht-RNAseq and (B) tar-RNAseq. In both heatmaps, the genes (rows) were sorted by median TPM values from low to high, and the samples (columns) were sorted by RIN from low to high.

**Supplementary Figure 6.** The coefficient of variation for relative expression (TPM) was lower for tar-RNAseq (green) than for wth-RNAseq) (orange) in samples with low RNA integrity (RIN < 3) and similar in samples of RIN > 3.
